# Interpretable antibody-antigen interaction prediction by introducing route and priors guidance

**DOI:** 10.1101/2024.03.09.584264

**Authors:** Yutian Liu, Zhiwei Nie, Jie Chen, Xinhao Zheng, Jie Fu, Zhihong Liu, Xudong Liu, Fan Xu, Xiansong Huang, Wen-Bin Zhang, Siwei Ma, Wen Gao, Yonghong Tian

## Abstract

With the application of personalized and precision medicine, more precise and efficient antibody drug development technology is urgently needed. Identification of antibody-antigen interactions is crucial to antibody engineering. The time-consuming and expensive nature of wet-lab experiments calls for efficient computational methods. Taking into account the non-overlapping advantage of current structure-dependent and sequence-only computational methods, we propose an interpretable antibody-antigen interaction prediction method, S3AI. The introduction of structural knowledge, combined with explicit modeling of chemical rules, establishes a ‘sequence-to-function’ route in S3AI, thereby facilitating its perception of intricate molecular interactions through providing route and priors guidance. S3AI significantly and comprehensively outperforms the state-of-the-art models and exhibits excellent generalization when predicting unknown antibody-antigen pairs, surpassing specialized prediction methods designed for out-of-distribution generalization in fair comparisons. More importantly, S3AI captures the universal pattern of antibody-antigen interactions, which not only identifies the CDRs responsible for specific binding to the antigen but also unearths the importance of CDR-H3 for the interaction. Structure-free design and superior performance make S3AI ideal for large-scale, parallelized antibody optimization and screening, enabling the rapid and precise identification of promising candidates within the extensive antibody space.

## 1 Introduction

As the important immune molecules of the human immune system [1], antibodies are a specialized type of protein with the primary role of recognizing and combating invading pathogens [2, 3]. The interaction between antibodies and antigens is characterized by a remarkable specificity and plays a pivotal role in this immunological process [3]. Therefore, humans continue to explore methods for preparing antibodies to develop antibody drugs and try to apply them in clinical treatments [4, 5]. In recent years, the research and development of antibody drugs have entered a new stage, driven by the continuous development of technologies such as genomics [6], proteomics [7], and immunology [8]. Currently, antibody design methods with higher precision and efficiency are urgently needed to meet the promotion and application of personalized treatment and precision medicine [9, 10].

The core of improving the design efficiency of antibody drugs lies in estimating the interaction strength of antibodies and antigens. The interaction strength between an antibody and an antigen extends beyond a mere binary interaction; it should be characterized as a continuous variable. Quantified by metrics such as the dissociation constant (*K*_*d*_) or the half-maximal inhibitory concentration (*IC*_50_), this continuum encapsulates the depth and strength of the interaction. Although there have been several conventional wet-lab experiments [11–14] to estimate the interactions between antibodies and antigens, they remain expensive and time-consuming. This presents a significant barrier to thoroughly exploring the extensive antibody optimization space, limiting the identification of novel and potentially more effective antibody candidates. Therefore, efficient computational methods are urgently needed to predict antibody-antigen interaction (AAI) to promote antibody optimization and screening.

As shown in Fig.1a, current computational methods, especially deep-learning methods, can be mainly divided into structure-dependent and sequence-only methods for characterizing Antibody-Antigen Interactions (AAI). The former exhibits lower task complexity since the model takes protein tertiary structure as input, which is closely related to protein interactions [15–19]. Srivamshi et al. [15] employed graph convolutional networks and an attention layer to explicitly encode the partner’s context in an antibody-antigen complex. While PInet [16] encoded proteins as surface point clouds with physicochemical properties and 3D geometry, predicting epitope-paratope in antibody-antigen. Another line of research is redesigning the complementarity-determining regions (CDRs) to get a broad-spectrum antibody. Shan et al. [20] developed a geometric neural network with attention mechanisms for antibody’s CDR sequences optimization. However, experimental structure determination methods, such as NMR spectroscopy and X-ray crystallography, prove to be both time-intensive and expensive, resulting in limited available high-quality structures of antibodies and their complexes. Although protein structure prediction methods [21–23] have made break-through progress, their inherent errors [24, 25] and the structural characteristics of antibodies [26, 27] bring obvious interference to AAI prediction. Naturally, high-quality data insufficiency presents an obstacle to training deep-learning models with ideal generalization capability. In contrast, sequence-only methods, which take advantage of extensive antibody sequence data, offer a more efficient framework for large-scale antibody screening [28–33]. Earlier studies, such as ProABC [28], utilized a sequence-only random forest algorithm to predict paratope residues, eliminating the need for structured data to achieve accurate predictions. Parapred [29] employed a combination of local and global features, harnessing deep learning techniques to analyze AAI. Marson et al. [30] introduced a convolutional neural network (MCNN) to predict the antigen specificity of antibodies. DeepAAI [31] proposes dividing antigens and antibodies into ‘seen’ and ‘unseen’ collections, utilizing graph neural networks to model the issue of out-of-distribution (OOD) data. However, a common drawback shared by these sequence-only methods is their limited interpretability and prediction performance, primarily stemming from the alienation of one-dimensional sequences from functionality.

**Fig. 1.**
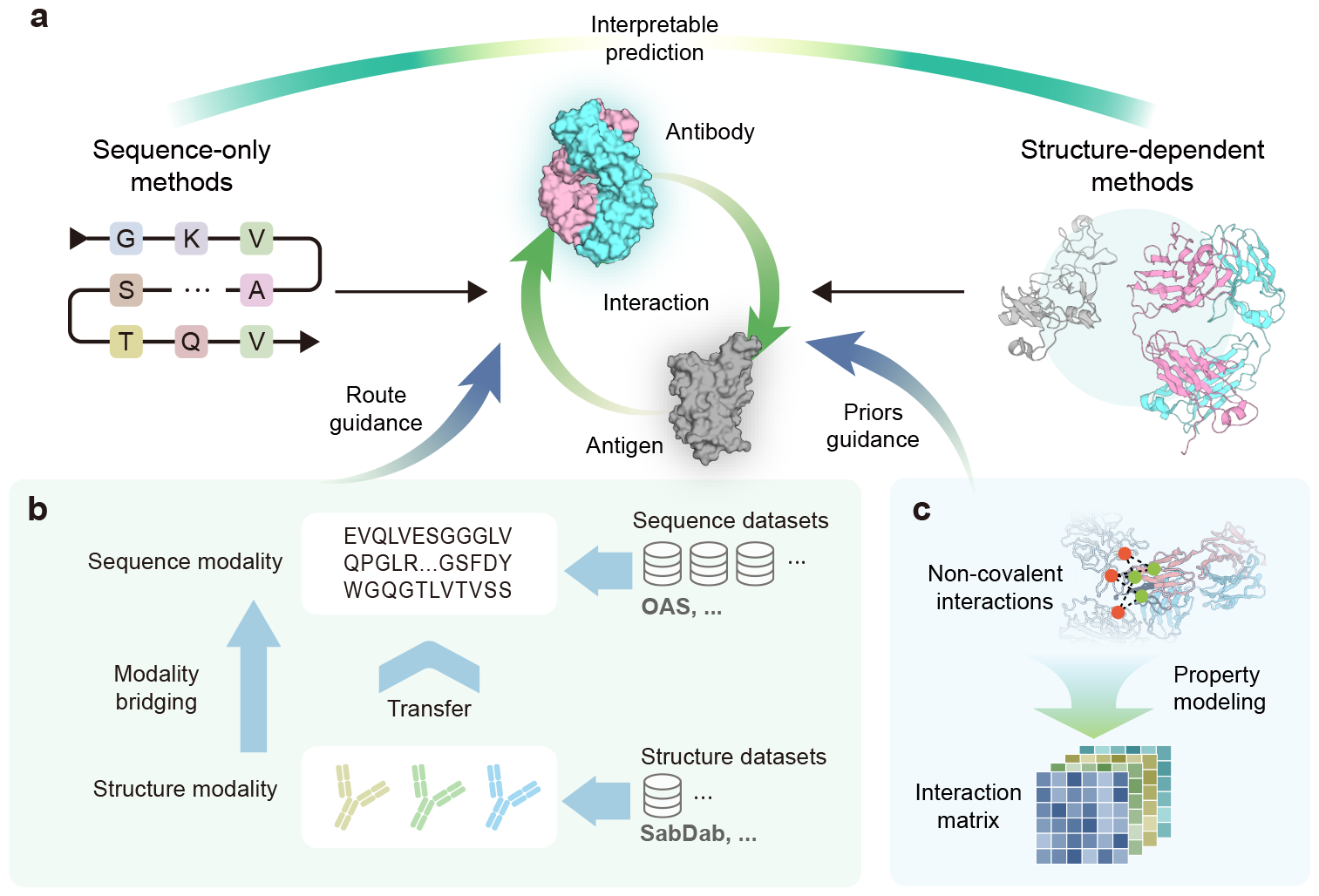
Our motivation and methodology. Route and priors guidance are designed to facilitate interpretable prediction of antibody-antigen interactions. **a**, Two major categories of methods currently used for antibody-antigen interaction prediction are sequence-only and structure-dependent methods. **b**, Structural information distillation transfers structural information into sequence modality through modality bridging, thereby providing route guidance for interpretable prediction. **c**, Interaction matrix makes non-covalent interactions explicit through property modeling, thereby providing priors guidance for interpretable prediction.

In order to combine the advantages of both types of methods while avoiding the disadvantages, it is essential to provide sensible ‘sequence-to-function’ guidance to structure-free models. Thus, we propose S3AI (**S**tructure-**A**ssisted **A**ntibody-**A**ntigen **I**nteraction Prediction), a deep-learning method that uses only sequences as inputs to the inference phase.

The first form of guidance we introduce aims to establish an implicit route from sequence to structure modalities. Modality bridging is a tangible way to build this route (Fig.1b), where we automatically incorporate protein structural information through a structural encoding module. By promoting the model’s ability to map from sequences to structural latent codes via contrastive learning, we break the barrier between these distinct modalities. In this process, knowledge from the 3D structures of massive antibodies is captured in the model parameters, allowing the model to introduce implicit structural information automatically in case only sequence input is used. The S3AI model incorporates two sequence feature extraction modules: one dedicated to antigens and the other to antibodies. Considering the volume of available structure and sequence data, structural information distillation is applied exclusively to the antibody feature extraction branch. Subsequently, the derived parameters are fine-tuned in several downstream tasks, aiming to enhance the performance of AAI predictions.

Another pivotal guidance of S3AI is the explicit modeling of chemical priors (Fig.1c). Recognizing the crucial role of non-covalent interactions in antibody-antigen binding, we devise a module to capture chemical constraints accurately throughout antigen-antibody docking. Beyond the features extracted from sequences using the protein language model, we generate various property maps of non-covalent interactions from the sequences, adhering to universally applicable chemical rules. This provides the model with insights into potential interaction formations through the integration of chemical priors in the deep-learning framework. The concatenated interaction matrix is passed through convolutional layers, facilitating effective feature aggregation and providing the model with a rich source of localized information. This novel module not only improves interpretability by affording a tangible linkage between molecular interactions and predictive outcomes but also provides local features that synergize with global features extracted in other stages.

Overall, S3AI is an interpretable deep-learning method that introduces route and priors guidance towards understanding antibody-antigen interaction. Downstream tasks for SARS-CoV-2 and HIV demonstrate the exceptional proficiency of S3AI in predicting AAI, surpassing current state-of-the-art predictors. Importantly, S3AI captures universal patterns of antibody-antigen interactions, revealing antigen-specific binding mechanisms and highlighting sub-regions important for the interaction. Moreover, S3AI’s capability to operate without requiring structural input during the inference phase positions it as an ideal solution for large-scale antibody optimization and screening. Naturally, S3AI can be applied to the design of antibodies for various targets, making the development of antibody drugs more precise and personalized.

## 2 Results

### 2.1 S3AI harnesses the property-driven architecture that bridges structure to sequence

S3AI achieves unparalleled accuracy and throughput in predicting antibody-antigen interactions by effectively integrating structural information with sequence input. Prior to the advent of S3AI, methods focusing solely on sequence inputs attempted to predict metrics related to AAI directly from the sequences of antibody-antigen pairs. However, these approaches often fell short due to their lack of structural information, which is crucial for understanding the nuances of interactions. Consequently, the ‘routes’ constructed are typically blind, intricate, and lack interpretability, making it challenging to derive meaningful insights or predictions.

As shown in Fig.2a, S3AI, on the other hand, revolutionizes this approach by actively guiding the mapping from antibody sequences to their structures, thereby providing a more rational and coherent route for predicting AAI from sequence inputs. The above strategy is here called ‘structural information distillation’, and its core lies in training a structural encoder through contrastive learning to obtain structure-enhanced features. In the teacher network, the antibody structure is processed by a pre-trained structure network to extract structural features. In the student network, the antibody sequence is fed to the protein language model (*i*.*e*.ESM [34] and then encoded by a learnable structure encoder to obtain features that incorporate structural information. Finally, the training manner of contrastive learning allows the knowledge from the antibody structure to be transferred to the structure encoder. This manner not only leverages the inherent sequence information but also enriches it with structural data, enhancing the model’s performance and the interpretability of its predictions. By doing so, S3AI offers a more informed and precise framework for understanding the intricate dynamics of antibody-antigen interactions.

**Fig. 2.**
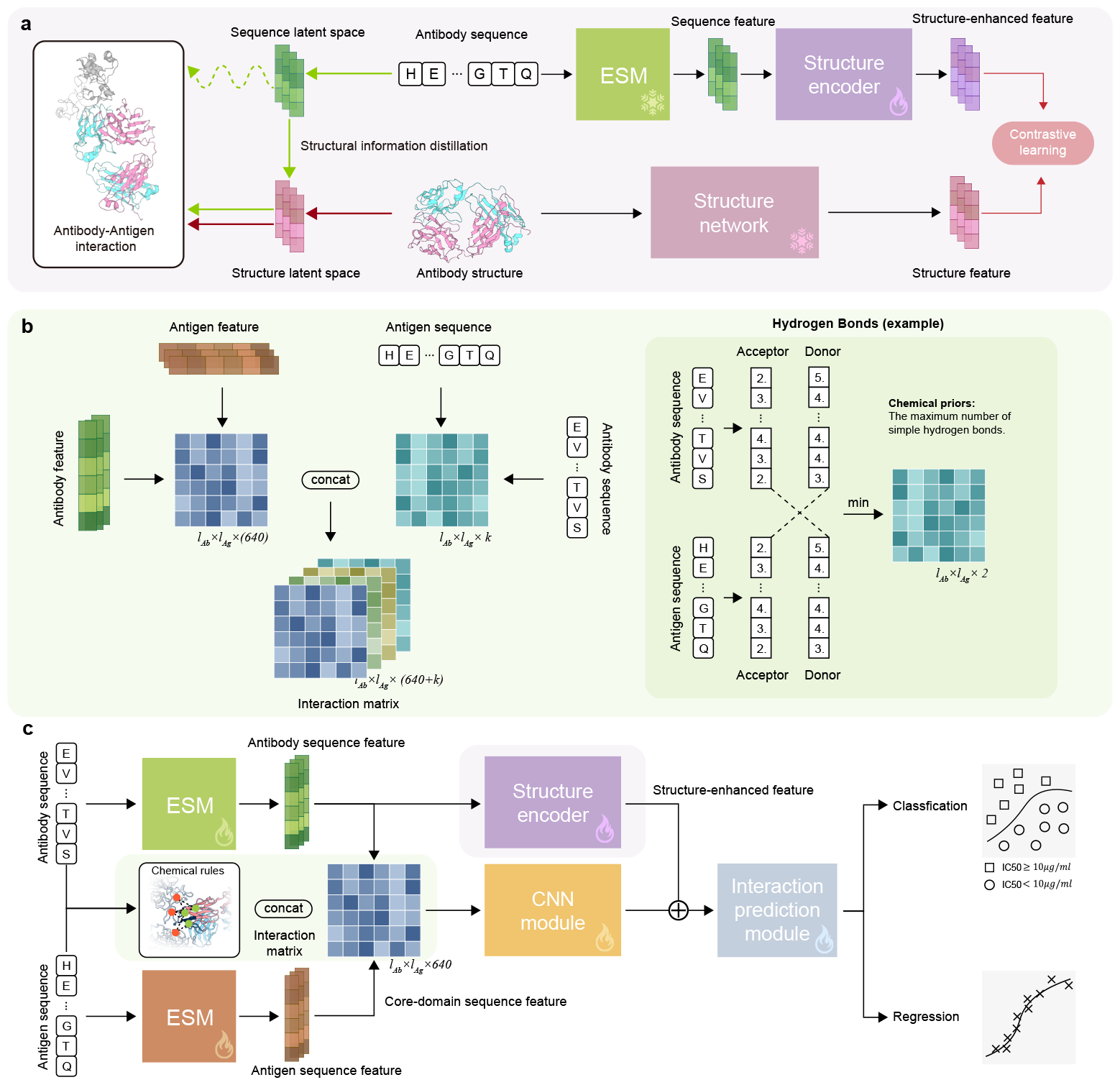
**a**, Structural information distillation, in which the antibody structure information extracted by the structure network is distilled into antibody sequence modality, thereby training the structure encoder to obtain structure-enhanced features. **b**, Interaction matrix, in which the feature matrix is calculated by implicit sequence embeddings, and the property matrix contains the physicochemical properties calculated by chemical rules. **c**, The overall framework of S3AI. The input antibody sequence is extracted with structure-enhanced feature, and the interaction matrix is concatenated with it after passing through the CNN module. The concatenated feature is then input into the subsequent interaction prediction module for classification (neutralization or not) and regression (*IC*_50_ estimation) tasks.

Implicit modeling of AAI based on deep learning models is challenging because of the complexity of molecular interactions, which can result in fundamental chemical rules being poorly understood by the neural network. To address this challenge, we propose the interaction matrix, which explicitly models chemical rules to introduce crucial non-covalent interactions. As shown in Fig.2b, the sequences of an antibody and an antigen are taken to calculate a property matrix containing non-covalent interactions, including hydrogen bonding, electrostatic interactions, van der Waals forces, and hydrophobic interactions. Take hydrogen bonding as an example, to construct a H-bond map between an antibody and an antigen, we initially analyze their sequences to identify the number of hydrogen bond donors and acceptors for each amino acid. This step produces two vectors respectively, with lengths matching those of the antibody and antigen, where each vector element indicates the count of hydrogen bond donors or acceptors at every amino acid position. Next, we compare the corresponding amino acids between the antibody and antigen, determining the minimum number of available donors and acceptors for each pair. This results in two matrices, sized by the antibody and antigen sequence lengths. These matrices delineate the potential H-bond interactions between each amino acid pair, pinpointing regions with high interaction propensity. Furthermore, the sequences of both the antibody and the antigen are processed to extract embeddings using protein language model (ESM here), and these embeddings are then used to create a feature matrix. The above property matrix and feature matrix are concatenated to form the final interaction matrix, which contains both implicit and explicit interaction patterns.

With the support of structural information distillation and interaction matrix, the overall architecture of S3AI is shown in Fig.2c. The antibody and antigen sequences are passed through the protein language model (ESM here) to obtain corresponding sequence features, in which the sequence features of core-domain of the antigen—determined by the antigen’s specific type—are adopted to form the feature matrix in the interaction matrix. The interaction matrix performs feature aggregation through the CNN module and is concatenated with the structure-enhanced feature of the antibody obtained through the structure encoder. The above-mentioned concatenated features are input into the interaction prediction module for downstream tasks in the manner of multi-task learning, including the binary classification task of whether the antibody-antigen pair is neutralizing and the regression task of *IC*_50_ estimation.

### 2.2 S3AI significantly outperforms state-of-the-art models

In order to comprehensively evaluate the predictive ability of S3AI for AAI, we collect SARS-CoV-2-related AAI data from previous studies and integrate them into the largest dataset to date. As shown in Fig.3a, we perform a thorough comparison with a series of models for antibody-antigen interaction prediction on regression tasks and classification tasks, including MCNN [30], Parapred [29], Fast-Parapred [35], AG-Fast-Parapred [35], PIPR [36], and ResPPI [37]. The detailed implementation can be found in Supplementary information, Section S1. For the regression task of *IC*_50_ estimation, S3AI significantly surpasses other models. It showcases superior performance by achieving a Spearman correlation coefficient of 0.655 and a Pearson correlation coefficient of 0.860. Compared to the best-performing existing model, this represents an improvement of approximately 13.3% in the Spearman correlation and about 7.1% in the Pearson correlation. For the classification task of neutralization, S3AI still comprehensively outperforms other models, with an accuracy of 84.53% and an MCC of 68.50%. Overall, S3AI shows comprehensive superiority in both regression and classification tasks, stamping a new watermark for antibody-antigen interaction prediction.

**Fig. 3.**
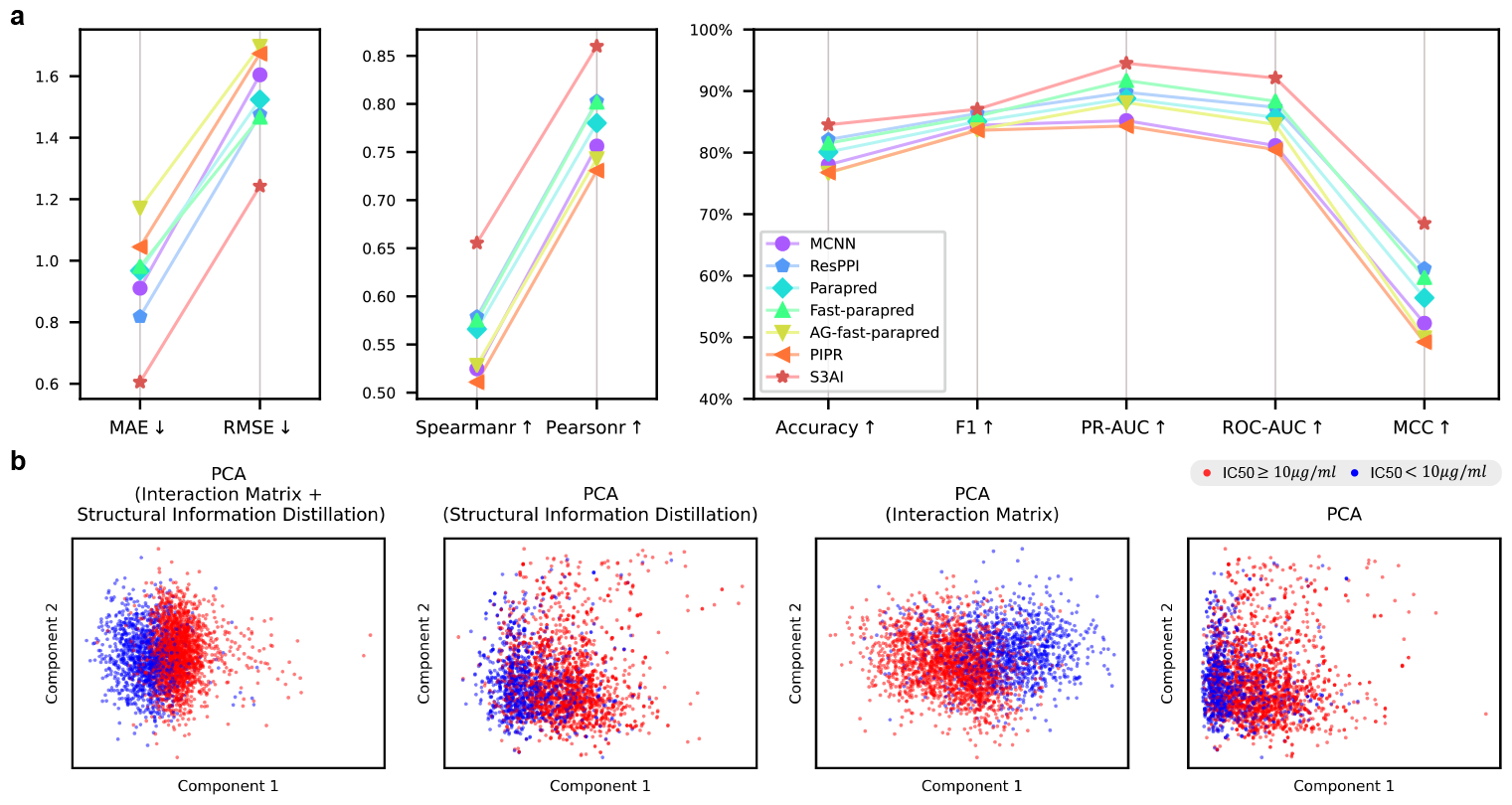
**a**, Prediction performance on integrated SARS-CoV-2 datasets. In both classification and regression tasks, S3AI outperforms state-of-the-art models across the board. **b**, Visualization of the impact of each module on the final concatenated features. The interaction matrix has a stronger impact on features than structural information distillation, and the joint use of the two modules achieves the best results.

Structural information distillation and interaction matrix are crucial to the prediction performance of S3AI, which is demonstrated in the visualization experiment in Fig.3b and Table.S4. Principal component analysis (PCA) is adopted to explore the impact of each module on the concatenated features, *i*.*e*., concatenations of structure-enhanced features, and CNN-processed interaction matrix. Compared with the visualization result on the far right without using the above two modules, the use of any module results in better clustering of features, and the joint use of the two modules achieves the best clustering effect. It is worth noting that the impact of the interaction matrix on the clustering effect is stronger than that of structural information distillation, which also confirms our hypothesis: it is difficult for neural networks to directly understand intricate molecular interactions, and explicit chemical priors modeling tends to provide a more accurate learning route.

### 2.3 S3AI’s generalization performance for out-of-distribution scenarios

Out-of-distribution (OOD) generalization is a common challenge faced by all deep learning models. The above challenge is even more pronounced in the prediction scenario of antibody-antigen interactions since most antibodies are ‘unseen’, *i*.*e*., the interactions of a large number of antibodies with any antigen are unknown [31]. For natural antibodies, the antibody space produced when faced with viral invasion is very large, which makes it time-consuming and costly to measure the interaction strength of any antibody-antigen pair through wet-lab experiments. In addition, the interactions of synthetic antibodies with antigens are also blind to us. Therefore, deep learning models for antibody-antigen interaction prediction require ideal OOD generalization capabilities.

Previous work proposed a deep-learning model, DeepAAI [31], specifically customized to predict interactions of ‘unseen’ antibody-antigen pairs. In order to evaluate the generalization performance of S3AI, we adopt the original architecture of S3AI to compare performance with DeepAAI without adding any customized modules for OOD. The only adjustment is that we change the multi-task framework of S3AI to the same single-task framework as DeepAAI for a fair comparison. As shown in Fig.4a, in the neutralization classification task, S3AI without customization completely surpassed all model variants of DeepAAI. On the regression task of *IC*_50_ estimation, S3AI surpasses most model variants, including the two model variants that only input sequences (Fig.4b). We can see that S3AI is slightly worse than the two model variants whose inputs contain position-specific scoring matrices (PSSMs) with evolutionary information. As an extremely time-consuming descriptor, PSSMs are calculated based on multiple sequence alignments and incorporate rich evolutionary information. However, S3AI is a deep-learning method that only requires input sequences and does not contain additional homologous information, which may explain why its prediction performance on regression tasks is slightly worse than that of the model variants using PSSMs of DeepAAI. Overall, S3AI still surpasses the state-of-the-art OOD model under the same input configuration without special modifications to adapt to OOD scenarios. This indicates that S3AI has learned universal patterns of molecular interactions and thus accurately predicts interactions of never-before-seen antibody-antigen pairs.

**Fig. 4.**
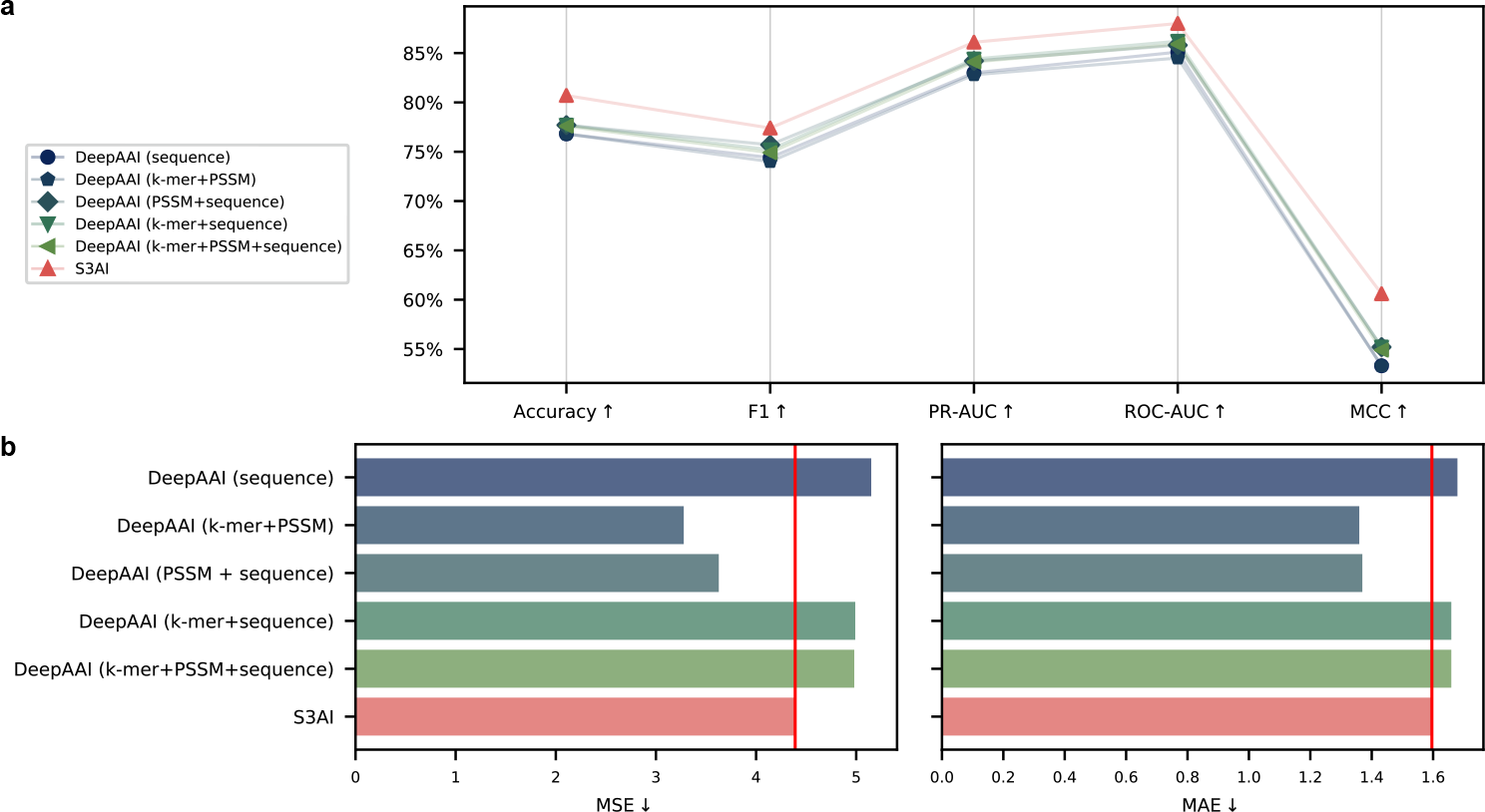
Generalization performance on HIV datasets. **a**, In the binary classification task of whether an antibody neutralizes HIV, S3AI comprehensively outperforms the state-of-the-art models specifically designed for out-of-distribution generalization of antibody-antigen interaction prediction, including model variants using PSSMs containing rich evolutionary information. **b**, In the regression task of *IC*_50_ estimation, S3AI outperforms most of the model variants, including all model variants that do not incorporate features with additional evolutionary information.

### 2.4 S3AI captures the universal pattern of antigen-antibody interactions

The aforementioned performance comparison demonstrates the excellent ability of S3AI in predicting AAI, and indicates that it has learned universal patterns of molecular interactions to a certain extent. What needs to be further explored is whether the interaction pattern extracted by S3AI is consistent with the actual antibody neutralization mechanism. To this end, we adopt a strategy of calculating the region effective attribution, in which the higher the attribution of a region to the prediction result, the more significant impact it has on the final prediction. There are basic modes of neutralization between different antibodies and different antigens, the core of which is that the Complementarity Determining Regions (CDRs) are responsible for binding to the antigen [38, 39]. As shown in Fig.5a, the region effective attributions of all antibody samples are displayed, and the CDRs of most samples have more significant attributions than other framework regions (FRs) in variable regions. The average attributions of all samples show that the impact of CDRs is several times that of FRs, which proves that CDR is the most important region for the interaction between antibody and antigen. In Fig.5b, the region effective attributions of four antibody samples are shown. The difference between these samples is that the attributions of FRs to the interactions have different characteristics. In sample 1519, all FRs are positive attributions, while in sample 5401, it is exactly the opposite, that is, all FRs are negative attributions. Furthermore, in sample 8485 and sample 27248, negative and positive attributions coexist. The above phenomenon confirms a basic fact that the attributions of FRs to the interactions are flexible and variable, which is caused by the binding characteristics of different antibody-antigen pairs.

**Fig. 5.**
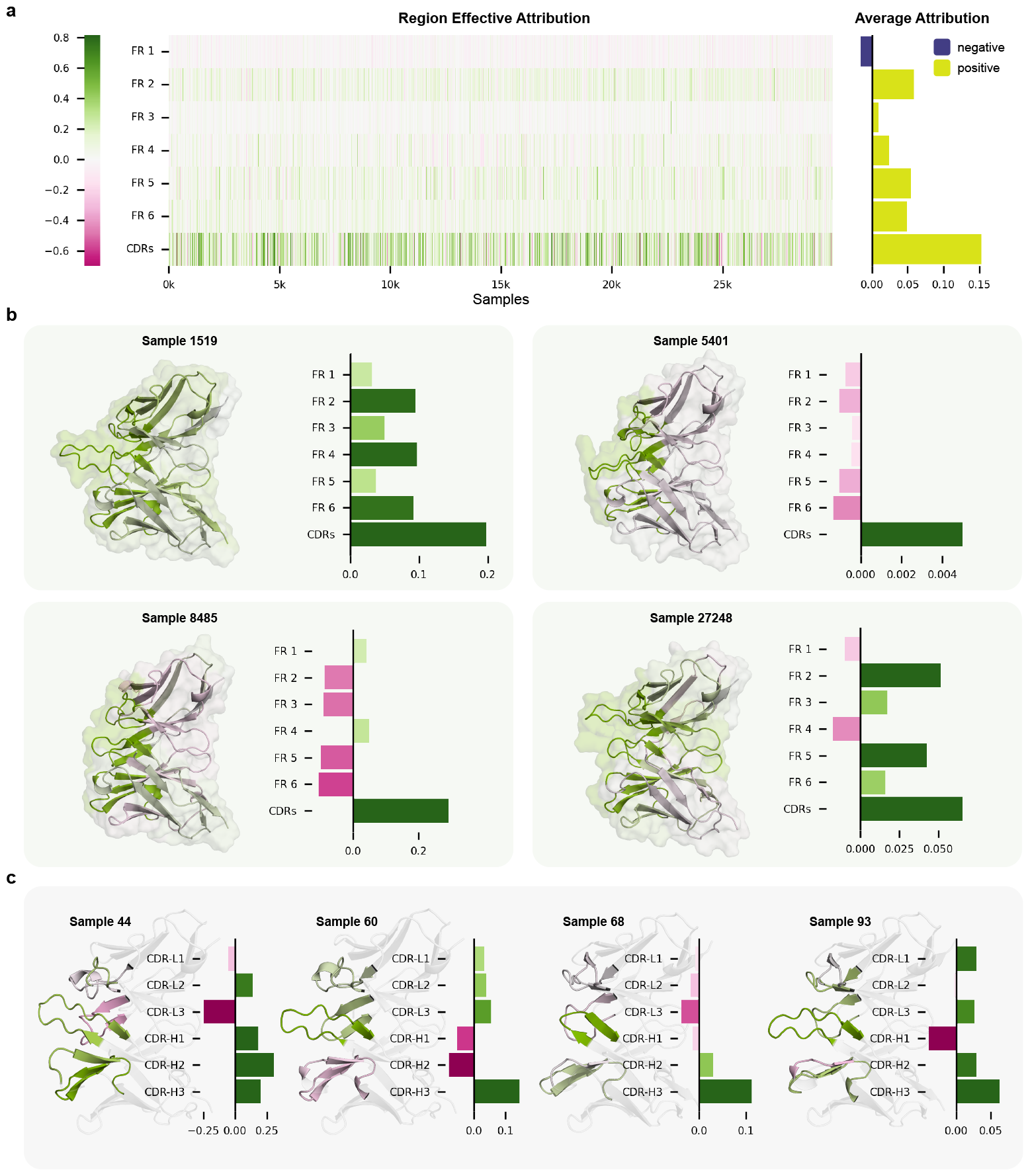
**a**, Effective attributions of each region in variable regions to the predictions of antibody-antigen interactions. Among them, CDRs have the most significant average attribution, which is consistent with the basic mode of antibody neutralization. **b**, Effective attributions of four samples with different characteristics. The structures of antibody variable regions are colored according to the degree of attributions. **c**, Analysis of effective attributions within CDR regions of four samples. Different sub-regions in the CDRs are colored according to their attributions. Among them, CDR-H3 always tends to have a significant effective attribution, indicating that this sub-region is crucial for antibody neutralization, which is consistent with previous research conclusions.

Moreover, CDR includes six sub-regions, namely CDR-L1, CDR-L2, CDR-L3, CDR-H1, CDR-H2, and CDR-H3. A large number of previous works have agreed that CDR-H3 is important for the interaction between antibodies and antigens, which makes us very curious whether S3AI can capture this phenomenon [40–44]. As shown in Fig.5c, sub-regions in the CDRs of four antibody samples are displayed. What the four samples have in common is that CDR-H3 always tends to have significant effective attribution, which is consistent with the above conclusion. Interestingly, the impacts of sub-regions within the CDRs, aside from CDR-H3, are diverse and irregular [41, 42, 45], which further highlights the complexity of AAI mechanism.

## 3 Discussion

Antibodies are important immune molecules that play a crucial role in the immune process. The remarkable specificity of antibodies against antigens has led humans to continue to explore technical routes to develop them into clinical drugs. With the increasingly urgent demand for personalized and precision medicine, the research and development of antibody drugs requires more efficient and high-precision technical means. Computational methods, especially deep learning methods, have shown great potential in antibody optimization or screening. However, the two current mainstream methods for predicting AAI, namely structure-dependent and sequence-only methods, have obvious shortcomings, which come from the scarcity of structural data and the alienation between sequence and function. In addition, the common difficulty faced by the above two types of methods is that it is challenging for neural networks to understand intricate molecular interactions. Therefore, the development of an interpretable deep-learning model for AAI prediction that overcomes structural or sequence mode constraints is of great significance for antibody engineering.

The first contribution of S3AI is to break the barrier between structure-dependent and sequence-only methods. Predicting AAI based on sequences has a high task complexity, which is rooted in the alienation route from sequence to structure and then to function. This motivates us to utilize structural information implicitly, that is, to incorporate knowledge from the protein structure modality to the sequence modality. In this work, we propose a strategy named structural information distillation to achieve modality bridging. First, a teacher network with structure as input and a student network with sequence as input are set up. The training manner of contrastive learning is further used to store the information from the teacher network to the structure encoder. The structural encoder is capable of transforming sequence features into structure-enhanced features, eliminating the need for input structure in the downstream task training and inference stages. In short, structural information distillation effectively realizes cross-modal information transfer and promotes the establishment of ‘sequence-to-function’ route.

S3AI’s second contribution is the explicit modeling of chemical priors to guide neural networks in understanding intricate molecular interactions. Antibody-antigen interactions are primarily driven by non-covalent interactions, so we propose interaction matrix to introduce the composite impact of hydrogen bonding, van der Waals forces, electrostatic interaction, and hydrophobic interactions. In general, this interaction matrix makes up for the shortcomings of the implicit modeling of AAI. In other words, explicit modeling of interaction patterns reduces the task complexity of predicting functions directly from sequences.

The design of the above two modules ensures S3AI’s excellent performance in predicting AAI. In this work, we comprehensively examine the capabilities of S3AI, including its generalization performance for out-of-distribution scenarios. S3AI not only significantly surpasses the state-of-the-art models but also exhibits ideal OOD generalization capabilities. Without any adjustments for the OOD task, S3AI still outperforms all variants of the state-of-the-art model customized for OOD generalization in predicting AAI in the classification task of neutralization or not. In the regression task of *IC*_50_ estimation, S3AI outperforms all model variants that only adopt sequences as input in a fair comparison. The above prediction performance indicates that S3AI has a certain degree of understanding of the basic patterns of molecular interactions, which facilitates its generalization for the prediction of unknown antibody-antigen pairs. We further intuitively explore the ability of S3AI to capture the universal pattern of AAI. Consistent with the antibody neutralization mechanism, S3AI accurately identifies CDRs responsible for specific binding to antigens. Furthermore, the importance of sub-region CDR-H3 for AAI is also discovered by S3AI. The above results demonstrate S3AI’s awareness of molecular interaction mechanisms, which is rooted in the introduction of route and priors guidance.

The potential of S3AI lies in the large-scale optimization and screening of antibodies. The structure-free input mode in the inference stage breaks through the dilemma of the scarcity of high-quality antibody structures, ensuring that antibody optimization and screening are only performed at the sequence level and can be massively parallel. This will greatly improve the efficiency of discovering immunologically active or specific antibodies in the huge antibody space, thereby accelerating the development of antibody drugs. However, the prediction performance of S3AI is limited by the low quality of distilled structural information, which is caused by the predicted antibody structures used in the structural information distillation module. It is foreseeable that in the future, as the scale of available high-quality antibody structures increases, the structural information distillation module will be able to extract more general and precise implicit structural information. In addition, further accurate modeling of chemical rules is also a feasible way to improve the prediction performance of S3AI. There is also an expectation for a lighter and faster prediction method that could further hasten antibody engineering.

## 4. Methods

### 4.1 Data

#### 4.1.1 Data for structural information distillation

The antibody structure data utilized for structural information distillation in this study encompasses two main types: real antibody data collected from the SabDab [46] and predicted antibody structure data derived from sequences using Igfold [47], which is a deep learning model specifically designed for predicting antibody structures. For the real antibody data, we exclude single-chain antibodies, entries with missing information, those with formatting errors (such as redefined atoms), and sequences with discontinuities within the chains. The structural data predicted by IgFold includes paired antibodies from the OAS dataset, along with a selection of relevant antibodies from the coronavirus.

#### 4.1.2 SARS-CoV-2 data

Given the absence of comprehensive studies summarizing *IC*_50_ data across various antibody-antigen pairs in the field, *IC*_50_ data are collected from several biological studies [48–51] that measured and published *IC*_50_ values between SARS-CoV-2 or its variants and human antibodies. These studies encompassed 18 different lineages, including those that emerged after Omicron. We obtain mutations of each lineage from outbreak.info [52, 53]. These mutations are then introduced into the spike protein sequences of the wild-type coronavirus, as obtained from GISAID [54]), thereby generating the spike protein sequences for each lineage.

Manual standardization is necessary because the *IC*_50_ values originated from diverse papers with varying experimental conditions. For example, the reporting of negative sample values (where *IC*_50_ *≥*10 *µ*g*/*ml) varies across studies, with some listing them as ‘*>*10’, ‘*>*100’, or specifying a particular value over 10, or even a constant value (*e*.*g*., 1000 *µ*g*/*ml). In our study, we standardize all negative *IC*_50_ samples to a uniform value of 10 *µ*g*/*ml. Similarly, we classify samples with *IC*_50_ *≥* 10 *µ*g*/*ml as non-neutralizing, while those with values below 10 *µ*g*/*ml are deemed neutralizing.

Moreover, during the de-duplication process of antibody sequences, discrepancies in *IC*_50_ values for the same antibody-antigen pair across different studies are observed. Such inconsistent data are identified as errors and excluded to ensure dataset accuracy. We yield *IC*_50_ values for 29,483 pairs of coronavirus antigens and corresponding antibodies.

#### 4.1.3 HIV data

The HIV data used to test OOD performance in our study comes from the Compile Analyze and Tally NAb Panels (CATNAP) at the Los Alamos HIV Database (LANL) [55], as published by DeepAAI. We follow the dataset’s split between ‘seen’ and ‘unseen’ data, conducting OOD tests on the ‘unseen’ dataset.

### 4.2 Architecture overview

S3AI is built on the protein language model that takes paired antibody and antigen sequences as input. The heavy and light chain sequences of the input antibody are concatenated and then encoded by ESM. For antigens, the protein sequence is fed into a separate ESM to produce the sequence feature:

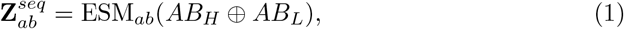

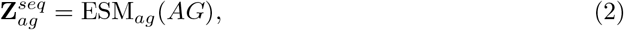

where *AB*_*H*_ = (*ab*_1_, *ab*_2_ *…, ab*_*k*_) and *AB*_*L*_ = (*ab*_*k*+1_, *ab*_*k*+2_, *…, ab*_*m*_) represent the sequences of the heavy and light chains of antibody, respectively, while *AG* = (*ag*_1_, *ag*_2_,, *ag*_*n*_) denotes the sequence of antigen. Structure encoder introduces structural information to sequence features of antibody 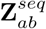, producing 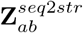 (see Section 4.3). Meanwhile, the sequence features 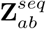, along with their respective sequences, are leveraged to generate the interaction matrix 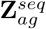 . This matrix is subsequently processed by Convolutional Neural Network (CNN) module, aimed at extracting interaction-related feature 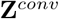 (refer to Section 4.4). Two different multilayer perceptrons (MLPs) are employed to subsequently process the concatenated features, yielding predictions for both classification and regression tasks:

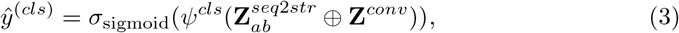

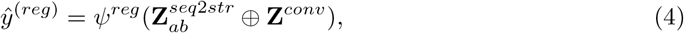

where *ψ*^*cls*^ and *ψ*^*reg*^ represent the MLPs for classification and regression tasks, respectively; *⊕* donates the concatenation operation on the final features.

### 4.3 Structural information distillation

The tertiary structure and sequence of antibodies can be considered two distinct modalities. In AAI problems, the structural modality often contributes more significantly than the sequence modality. However, the scarcity or even absence of structural data in many application scenarios leads to performance loss. To address this challenge, we look towards a series of cross-modal distillation methods [56–58], which offer a strategy to address the issue of missing modalities.

Building on this foundation and inspired by the recent advancements in the field of molecular property prediction, notably the 3DInfomax [59], we leverage contrastive learning to facilitate knowledge transfer between structural and sequence networks. This process can be regarded as a form of cross-modal distillation technique.

Structural information distillation involves two networks: a teacher network *f*_*str*_() that receives antibody structural inputs and outputs structural representations, and a student network *f*_*seq*_() that takes sequence inputs to generate sequence features. The teacher network’s weights are derived from pretraining on a protein structure dataset and remain fixed throughout this process. The student network, comprised of concatenated ESM and structure encoder, is tasked with extracting features from the antibody sequence. During the training process, the student network’s parameters are updated, enabling it to learn how to incorporate structural information into sequence features. For an antibody *x* = (*x*^*seq*^, *x*^*str*^) in a dataset, its sequence and structure are input into *f*_*seq*_() and *f*_*str*_(), respectively, yielding representations *z*^*seq*2*str*^ and *z*^*str*^. The training objective at this stage is to maximize the similarity between representations of the same antibody 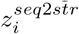 and 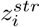 while minimizing the similarity between unmatched representations 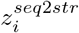 and 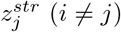 .

The similarity measurement function is defined as the cosine similarity, which is given by the formula:

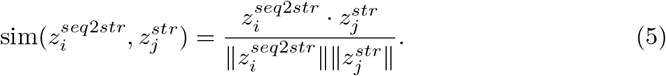

To guide this process, the NT-Xent (normalized temperature-scaled cross entropy) loss is utilized[60]:

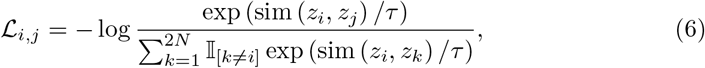

where the indicator function 𝕀 _[*k?*≠ *i*]_ takes values from {0, 1}, is used to determine whether *k* is not equal to *i* (evaluating to 1 if and only if *k* ≠ *i*). Additionally, *τ* represents a temperature parameter that adjusts the scale of the loss. The NT-Xent loss is calculated for all positive pairs within a mini-batch, ensuring symmetry in the evaluation of pairwise similarities.

We adopt the structural representation network proposed in [61] as our teacher network, utilizing the pretrained weights they provided. The teacher network is solely engaged in the process of structural information distillation, extracting representations from structural data, and does not participate in the training or inference processes of downstream tasks. The parameters of the structure encoder within the student network are used for initialization and undergo further fine-tuning on downstream tasks.

### 4.4 Interaction matrix

Within the array of structure-independent methods, beyond the previously mentioned challenge of diminished accuracy stemming from an absence of structural data, lies an additional complication: the issue of ambiguous interpretability. In response, several studies [62–65] have investigated the incorporation of attention mechanisms to enhance interpretability. However, the intricate nature of molecular interactions presents a significant challenge to the implicit modeling efforts of AAI through deep learning models.Thus, given two sequences, the antibody *AB* = (*ab*_1_, *ab*_2_ *…, ab*_*m*_) and the antigen *AG* = (*ag*_1_, *ag*_2_, *…, ag*_*n*_), we propose interaction matrix **Z**^*inter*^*∈* ℝ*m×n×d* to capture the interaction pattern between antibody and antigen. The interaction matrix is composed of two parts:

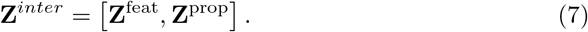

The feature matrix **Z**^feat^ is derived from the element-wise multiplication of latent codes, capturing sequence information:

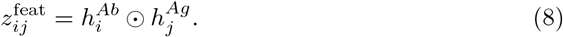

Here, 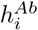 and 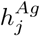 are the latent codes for the *i*^*th*^ amino acid in the antibody and the *j*^*th*^ amino acid in the antigen, respectively.

For each type of non-covalent interaction, the property matrix **Z**^prop^ introduces specific channels, including hydrogen bonding, electrostatic interactions, van der Waals forces, and hydrophobic interactions.

We first focus on hydrogen bonding, a fundamental interaction widely prevalent across various protein-protein interactions. For each pair of amino acids, *ab*_*i*_ from the antibody and *ag*_*j*_ from the antigen, the calculation for the H-bond matrix is as follows:

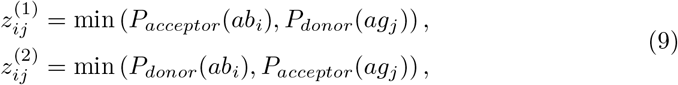

where *P*_*acceptor*_ and *P*_*donor*_ represent the counts of hydrogen acceptor and donor atoms, respectively, of an amino acid. This process iterates over each amino acid pair between the two sequences, generating a comprehensive interaction profile that captures potential hydrogen bonding.

Another type of important non-covalent interaction between antibodies and antigens is electrostatic interactions. These interactions are primarily governed by the charge properties of the amino acids. To incorporate electrostatic interactions into the property matrix, a specific channel is introduced, which quantifies the potential for electrostatic interactions between each pair of amino acids. The calculation for this part of the property matrix can be represented as follows:

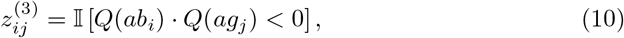

where *Q*(*x*) denotes the charge of amino acid *x*, which can be positive, negative, or neutral. This binary representation streamlines the modeling of electrostatic interactions by directly identifying when opposite charges are present between amino acids, thus indicating potential attractive forces that can enhance the interaction. In this context, only pairs of amino acids with opposite charges are considered to possibly contribute to the interaction potential.

Building on the foundation laid by the analysis of hydrogen bonding and electrostatic interactions, we next turn our attention to van der Waals forces. While subtler than the previously mentioned forces, van der Waals interactions play a crucial role in the nuanced dance of molecular recognition, acting as the fine threads that help weave the intricate tapestry of antibody-antigen interactions.

To capture potential van der Waals forces within interaction matrix, we introduce an additional layer of analysis. Based on the relationship between distances and interaction strengths between amino acids [66], we employ a Gaussian formula to approximate the effect of distance on van der Waals forces, taking into account the complementary nature of steric hindrance of amino acid side chains. The calculation of this addition to our property matrix is as follows:

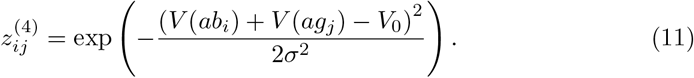

Here, *V* (*x*) denotes the van der Waals volume [67] of an amino acid *x. V*_0_ and *σ*^2^ are derived from a Gaussian fit to the aggregate of possible amino acid pair volumes.

As the final piece of our interaction matrix, we leverage the hydropathy index [68] to capture hydrophobic interactions. The calculation method is simple yet effective:

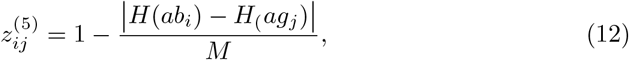

where *H*(*x*) represents the hydropathy index of amino acid *x*. The term *M* stands for the maximum absolute difference in hydropathy indices across all possible pairs of amino acids, ensuring the subtraction operation yields a normalized score that reflects the relative hydrophobic compatibility between *ab*_*i*_ and *ag*_*j*_. This channel provides information on hydrophobic interactions through a simple assessment of the differences in hydrophobic and hydrophilic between interacting amino acids.

Given the approximations and simplifications involved in creating the property matrix, as well as the complexity of actual interactions, we further extract features from the interaction matrix using a CNN module. This module enriches the network with local feature information. Through interpretability analysis (see Section 4.8), we have also proven its effectiveness in capturing the patterns of antibody-antigen interactions.

### 4.5 Multi-task focal loss

Due to the specificity of the interaction between antibodies and antigens, most of the recorded antibody-antigen pairs do not exhibit neutralization. This has resulted in an imbalance of positive and negative samples in our collected SARS-CoV-2 dataset. Additionally, differences in fitness among different virus lineages lead to a non-uniform data distribution at the lineage level. (see Supplementary information, Fig.S1) Moreover, since the *IC*_50_ values in the dataset come from different sources, and the experimental conditions and methods cannot be guaranteed to be completely consistent, the dataset also contains some unavoidable noise. To address these issues, we have adopted a multi-task framework (see Section 4.2) along with the corresponding focal loss [69, 70] for training downstream tasks. The overall loss consists of regression and classification loss:

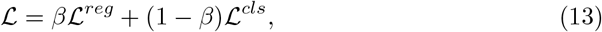

where *β* is a weighting factor to balance the regression and classification tasks. The regression part of focal loss is defined as:

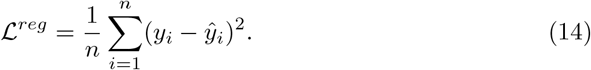

For the binary classification neutralization prediction task, we adopt the classical focal loss:

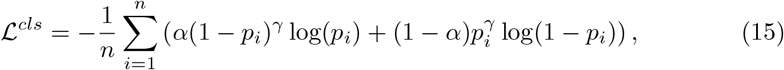

where (1 *− p*_*i*_) and *p*_*i*_ are the model’s estimated probabilities for the class with label *y* = 0 and *y* = 1, respectively; *α* is a weighting factor for the class; *γ* is the focusing parameter that smoothly adjusts the rate at which easy examples are down-weighted. (Supplementary information, Table.S8)

### 4.6 Evaluation metrics for regression and classification tasks

#### Mean Absolute Error (MAE)

MAE, conversely, measures the average magnitude of errors in a set of predictions without considering their direction. It’s calculated as the average of the absolute differences between predicted and actual values, thus providing an intuitive measure of prediction accuracy:

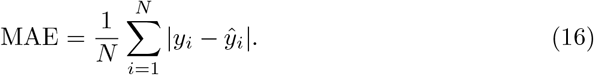

#### Root Mean Square Error (RMSE)

RMSE is a standard way to measure the error of a model in predicting quantitative data. The formula for RMSE is given by:

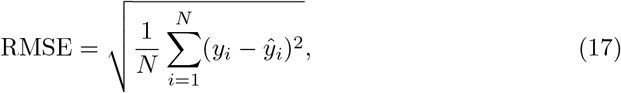

where *N* is the number of observations, *y*_*i*_ is the actual value of an observation, and 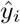 is the predicted value.

#### Pearson and Spearman Correlation

*IC*_50_ values originate from wet-lab experiments across various batches and different laboratories. Thus, given the diversity in experimental conditions, Spearman’s rank correlation coefficient (Spearman’s *ρ*) is particularly apt for measuring the relationship between ordinal variables, such as comparing the magnitudes of different *IC*_50_ values. It assesses how well the relationship between two variables can be described using a monotonic function, focusing on the ranks of values rather than their direct magnitudes:

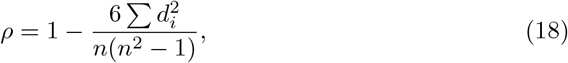

where *d*_*i*_ is the difference between the ranks of corresponding values, and *n* is the number of observations.

We also consider Pearson’s correlation coefficient (Pearson’s r) for its ability to measure the linear correlation between two variables, offering a comprehensive analysis when the data distribution permits:

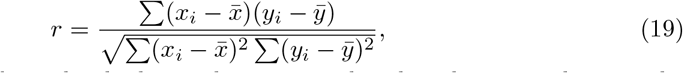

where *x*_*i*_ and *y*_*i*_ are the individual sample points indexed with *i*, 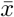 and 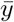 are the sample means.

Considering the variability in *IC*_50_ data due to different experimental setups, Spearman’s *ρ* becomes a more suitable choice for evaluating the ordinal relationship between *IC*_50_ values, providing a robust measure against the non-uniformity of data collection methods. Pearson’s r complements this by quantifying the degree of linear relationship where applicable, together offering a nuanced approach to assess predictive accuracy in the context of *IC*_50_ value prediction.

#### Accuracy, F1 Score, and Matthews Correlation Coefficient (MCC)

In evaluating binary classification models, especially with imbalanced datasets like the SARS-CoV-2 datasets, accuracy alone can be misleading. Thus, we supplement it with the F1 Score and the Matthews Correlation Coefficient (MCC). The F1 Score, calculated as the harmonic mean of precision *p* (correct positive predictions out of all positive predictions) and recall *r* (correct positive predictions out of all actual positives), offers a balanced metric:

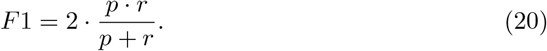

MCC further provides a comprehensive measure by accounting for all aspects of the confusion matrix, capturing the quality of binary classifications beyond the limitations of accuracy and F1 Score. This approach ensures a more accurate assessment of model performance in handling the skewed class distribution typical of the SARS-CoV-2 dataset.

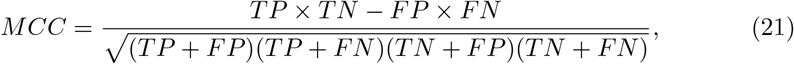

where TP is the number of true positives, TN is the number of true negatives, FP is the number of false positives, and FN is the number of false negatives.

#### Precision–Recall Area Under the Curve (PC-AUC) and Area Under the Receiver Operating Characteristic (ROC-AUC)

PR-AUC measures the trade-off between precision and recall across different thresholds. It is particularly valuable in the SARS-CoV-2 datasets, where positive samples (neutralizing pairs) are much less common than negative ones.

ROC-AUC evaluates how well the model distinguishes between neutralizing and non-neutralizing pairs over various threshold settings. It plots the true positive rate (recall) against the false positive rate (the ratio of incorrectly identified negatives) to show the model’s discrimination capability.

### 4.7 Module ablation study and principal component analysis

To further investigate and analyze the impact of structural information distillation and interaction matrix on model performance, we conduct module ablation study and PCA analysis on the final concatenated features.

We employ four configurations of the model: the original S3AI, one without structural information distillation, one without interaction matrix, and one without both components. For the version without structural information distillation, we randomly initialize the structural encoding module. For the variant lacking interaction matrix, we pool the sequence features of the antibody and antigen along their length, concatenate them with the other final features, and then feed them into the interaction prediction module for prediction. All models are trained under identical settings. After training, we use PCA analysis to visualize the concatenated features of the best checkpoint for each model on all samples in the test set. The final performance of ablation study is presented in Table.S4.

### 4.8 Region effective attribution calculation

#### Attribution evaluation

We use the Shapley value [71] to measure the attribution of the regions in the input antibody to the prediction result. The Shapley value is a renowned game-theoretic metric for assessing the attribution/importance of each input variable to the output of the deep learning model. It has been recognized as the sole attribution method that adheres to the axioms of anonymity, symmetry, dummy, additivity, and efficiency. Accordingly, the Shapley value [72] of the *i*-th variable is computed as follows:

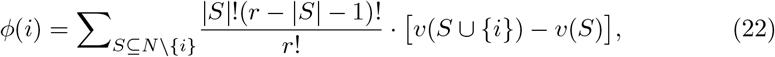

where N denotes the set of variables and | *·* | denotes the cardinality of the set. Here, we use *v*(*S*) to simplify the notation of *v*(*x*_*S*_), and *v*(*x*_*S*_) denotes the model output on a masked sample *x*_*S*_. In the masked sample *x*_*S*_, variables in *S* are present, and variables in *N \ S* are masked. In this way, *v*(*∅*) represents the model output when all input variables are masked, and *v*(*N*) denotes the model output on the original input sample *x*.

#### Attribution of FRs and entire CDRs

As the first step of estimating the attribution of the CDRs, we visualize the effects of the entire CDRs on the prediction result.

For an input antibody AB, let **Z**^*inter*^ *∈* ℝ^*m×n×d*^ denote the interaction matrix, where *m, n*, and *d* denote the length of the antibody AB, the length of the antigen AG and channel number of the interaction matrix, respectively. We select the feature of the entire CDRs as a single variable and divide the remained feature of FRs in **Z**^*inter*^ into *t* continuous regions. These regions and the CDRs are selected as *t* + 1 input variables, which consist of the input variable set *N*, to calculate the Shapley value. Then, we set *v*(*N*) as the prediction output for antibody-antigen interactions, and *v*(*S*) denotes the prediction output when the variable in *N \ S* is masked.

#### Attribution of each CDR sub-region

After calculating the attribution of the entire CDRs, we estimate the attribution of each CDR sub-region.

Given an intermediate-layer feature of an input antibody, the entire CDRs consist of 6 parts: CDR-H1, CDR-H2, CDR-H3, CDR-L1, CDR-L2, and CDR-L3. We select the feature of these 6 parts as input variables, which consist of the input variable set *N* . Then, we set *v*(*N*) as the attribution of the entire CDRs in this input antibody AB and *v*(*S*) denotes the attribution of the entire CDRs when the variable in *N \ S* is masked.

#### Mask method

When we calculate the prediction output *v*(*S*) on a masked input antibody, we systematically mask the selected variables in *N \S*, which represent features in the interaction matrix, and observe their impact on the model’s output. This method involves iteratively masking each feature while measuring the change in the model’s predictions. By comparing the model’s output with masked and present variables, Shapley values can be calculated to quantify the feature’s contribution to the prediction. This method requires a faithful baseline value to mask the variable, which does not provide additional prior information and does not cause the masked feature to be out of the distribution of the original features in the interaction matrix.

For the attribution of the entire CDRs, the baseline value to mask the variable is set as the variable’s average feature vector across the dataset, which is a widely used method to set the baseline value, providing a reference point for evaluating the significance of individual features.

For the attribution of each CDR sub-region, since *v*(*N*) is the attribution of the entire CDRs, we calculate the masked output *v*(*S*) by replacing the features of the regions in the CDRs that belong to *N\S* with baseline values when calculating the attribution entire CDRs. The baseline value to mask the variable is set as the variable’s average feature vector across the dataset.

#### Variable selection

In order to encourage greater faithful effective attribution, we generate many partitions of variables for calculating the Shapley value. In the selection of Shapley values for the attribution of the entire CDRs, the process involves randomly selecting contiguous regions from the input antibody after choosing CDRs as a whole, and for the attribution of each CDR sub-region, we also use these randomly selected regions to get several attribution value outputs.

Besides, these remaining regions can be randomly sampled in various sizes to create different scales of variables. These selected regions form the variables for computing the Shapley value. Through various combinations of these selections, attributions are calculated for each variable, and the average result across all selections represents the final attribution of the entire CDRs.

By calculating the attribution for each variable and averaging the results across all partitions, we obtain a comprehensive assessment of the entire CDRs’ attribution to the prediction outcome. Additionally, by averaging the attribution values outputs {*v*(*S*)}.from all partitions, this approach ensures a comprehensive evaluation of the attribution for each CDR sub-region.

## Data availability

The SabDab data is available at https://opig.stats.ox.ac.uk/webapps/sabdab-sabpred/sabdab. The processed HIV data is freely available at https://github.com/enai4bio/DeepAAI. SARS-CoV-2 data can be downloaded from https://github.com/stau-7001/S3AI.

## Code availability

Relevant code and models are available at https://github.com/stau-7001/S3AI.

## Acknowledgements

This work is financially supported by the National Key R&D Program of China (No. 2022ZD0118201, 2020YFA0908100, 2023YFF1204401), Natural Science Foundation of China (No. 61972217, 32071459, 62176249, 62006133, 62271465, 61825101, 62088102, 21991132, 21925102, 92056118, 22331003, 22301010), Shenzhen Medical Research Fund (No. B2302037), and Beijing National Laboratory for Molecular Sciences (BNLMS-CXXM-202006). The work is also supported by the National Center for Protein Sciences at Peking University. W.-B.Z. acknowledges Bayer Pharma for the Bayer Investigator Award. We appreciate the support of AI for Science (AI4S)-Preferred Program, Peking University Shenzhen Graduate School, China.

## Supplementary information

### S1 Baseline methods

In conducting the comparative experiments, as the models utilized for comparison are not all originally designed for *IC*_50_ regression or classification tasks, we implement several essential adjustments. For the Parapred-series models (Parapred, Fast-Parapred, and AG-Fast-Parapred), originally intended for predicting binding sites, as well as the PPI prediction models PIPR and ResPPI, we modify their output head to replicate the structure of S3AI, thereby enabling the prediction of *IC*_50_ values and binary classification outputs. For MCNN, in addition to the modifications mentioned above, we incorporate a module with the same architecture as the one originally designed for processing antibody inputs. This new module is designed for handling antigen inputs, allowing MCNN to accommodate the diverse antigen inputs in the dataset. All models are trained using the same multitask focal loss as employed by S3AI.

### S2 Overview of SARS-CoV-2 data

**Fig. S1.**
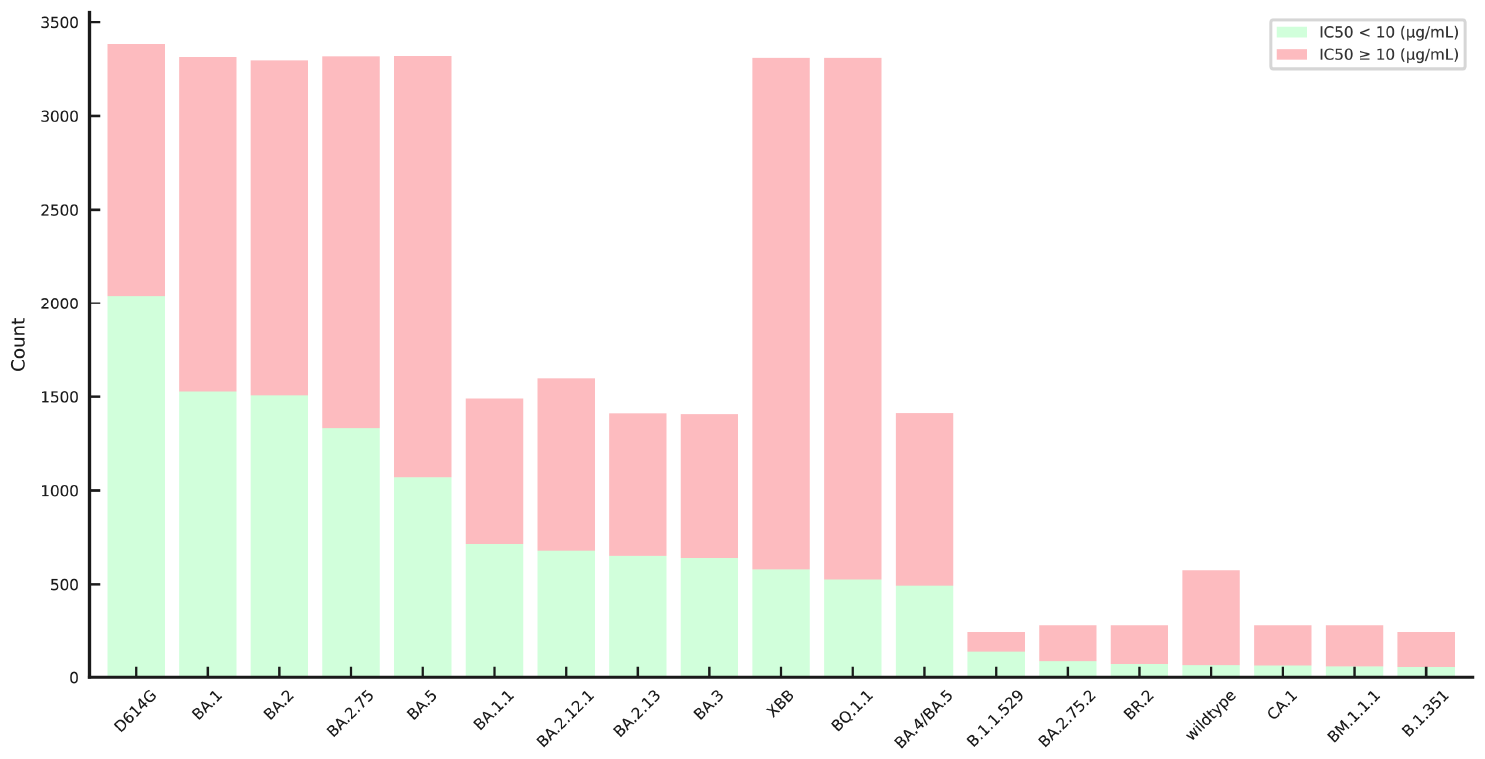
The distribution of *IC*_50_ values across different SARS-CoV-2 lineages

### S3 Results for SARS-CoV-2 test set

**Table S1.** Performance of models on SARS-CoV-2 test set.

| Metric | MCNN | ResPPI | Parapred | Fast-parapred | AG-fast-parapred | PIPR | S3AI |
| --- | --- | --- | --- | --- | --- | --- | --- |
| MAE ↓ | 0.911 | 0.818 | 0.967 | 0.981 | 1.171 | 1.045 | 0.605 |
| RMSE ↓ | 1.605 | 1.475 | 1.524 | 1.467 | 1.697 | 1.673 | 1.243 |
| Spearmanr ↑ | 0.525 | 0.579 | 0.566 | 0.575 | 0.528 | 0.511 | 0.655 |
| Pearsonr ↑ | 0.756 | 0.803 | 0.780 | 0.802 | 0.743 | 0.731 | 0.860 |
| Accuracy ↑ | 0.780 | 0.821 | 0.801 | 0.815 | 0.767 | 0.768 | 0.845 |
| F1 score ↑ | 0.844 | 0.863 | 0.851 | 0.859 | 0.837 | 0.836 | 0.870 |
| PR-AUC ↑ | 0.852 | 0.898 | 0.888 | 0.917 | 0.881 | 0.843 | 0.945 |
| ROC-AUC ↑ | 0.811 | 0.874 | 0.857 | 0.884 | 0.846 | 0.805 | 0.921 |
| MCC ↑ | 0.523 | 0.612 | 0.564 | 0.597 | 0.499 | 0.492 | 0.685 |

### S4 Results for HIV test set

**Table S2.** Classification Performance of models on OOD test (HIV unseen test set for classification).

| Model | Accuracy ↑ | F1 score ↑ | PR-AUC ↑ | ROC-AUC ↑ | MCC ↑ |
| --- | --- | --- | --- | --- | --- |
| DeepAAI (sequence) | 0.768 | 0.744 | 0.830 | 0.851 | 0.533 |
| DeepAAI (k-mer+PSSM) | 0.768 | 0.740 | 0.828 | 0.845 | 0.533 |
| DeepAAI (PSSM+sequence) | 0.777 | 0.757 | 0.842 | 0.858 | 0.552 |
| DeepAAI (k-mer+sequence) | 0.777 | 0.752 | 0.844 | 0.862 | 0.552 |
| DeepAAI (k-mer+PSSM+sequence) | 0.776 | 0.749 | 0.841 | 0.859 | 0.549 |
| S3AI | 0.807 | 0.774 | 0.861 | 0.880 | 0.606 |

**Table S3.** Regression Performance of models on OOD test (HIV unseen test set for regression).

| Model | MSE ↓ | MAE ↓ |
| --- | --- | --- |
| DeepAAI (sequence) | 5.15 | 1.68 |
| DeepAAI (k-mer+PSSM) | 3.28 | 1.36 |
| DeepAAI (PSSM + sequence) | 3.63 | 1.37 |
| DeepAAI (k-mer+sequence) | 4.99 | 1.66 |
| DeepAAI (k-mer+PSSM+sequence) | 4.98 | 1.66 |
| S3AI | 4.39 | 1.59 |

### S5 Results for module ablation study

**Table S4.** Results for module ablation study, including four configurations of the model: the original S3AI, one without structural information distillation (Interaction Matrix), one without interaction matrix (Structural Information Distillation), and one without both components (Baseline).

| Model | Baseline | Structural Information Distillation | Interaction Matrix | S3AI |
| --- | --- | --- | --- | --- |
| MAE ↓ | 0.955 | 0.899 | 0.645 | 0.605 |
| RMSE ↓ | 1.634 | 1.578 | 1.262 | 1.243 |
| Spearmanr ↑ | 0.544 | 0.552 | 0.626 | 0.655 |
| Pearsonr ↑ | 0.752 | 0.761 | 0.854 | 0.860 |
| Accuracy ↑ | 0.791 | 0.799 | 0.837 | 0.845 |
| F1 score ↑ | 0.838 | 0.847 | 0.866 | 0.870 |
| PR-AUC ↑ | 0.876 | 0.881 | 0.936 | 0.945 |
| ROC-AUC ↑ | 0.836 | 0.842 | 0.909 | 0.921 |
| MCC ↑ | 0.545 | 0.559 | 0.660 | 0.685 |

### S6 Region effective attribution

**Table S5.** Average region effective attribution of FRs and entire CDRs in Fig.5a

| FR1 | FR2 | FR3 | FR4 | FR5 | FR6 | CDRs |
| --- | --- | --- | --- | --- | --- | --- |
| 1.601e-02 | 5.857e-02 | 9.106e-03 | 9.721e-02 | 2.373e-02 | 4.919e-02 | 1.526e-01 |

**Table S6.** Region effective attribution of FRs and entire CDRs in Fig.5b

| Sample | FR1 | FR2 | FR3 | FR4 | FR5 | FR6 | CDRs |
| --- | --- | --- | --- | --- | --- | --- | --- |
| 1519 | 3.180e-02 | 9.545e-02 | 5.059e-02 | 9.721e-02 | 3.774e-02 | 9.212e-02 | 1.978e-01 |
| 5401 | -7.455e-04 | -1.046e-03 | -4.464e-04 | -4.709e-04 | -1.063e-03 | -1.361e-03 | 4.985e-03 |
| 8488 | 4.115e-02 | -8.567e-02 | -8.890e-02 | 5.021e-02 | -9.608e-02 | -1.039e-01 | 2.902e-01 |
| 27248 | -9.986e-03 | 5.151e-02 | 1.751e-02 | -1.747e-02 | 4.270e-02 | 1.611e-02 | 6.545e-02 |

**Table S7.** Region effective attribution of CDRs in Fig.5c

| Sample | CDR-L1 | CDR-L2 | CDR-L3 | CDR-H1 | CDR-H2 | CDR-H3 |
| --- | --- | --- | --- | --- | --- | --- |
| 44 | -9.326e-03 | -1.902e-02 | -3.844e-02 | -1.459e-02 | 3.083e-02 | 1.128e-01 |
| 60 | 2.932e-02 | -3.881e-03 | 2.634e-02 | -3.949e-02 | 2.932e-02 | 6.291e-02 |
| 68 | -5.323e-02 | 1.369e-01 | -2.485e-01 | 1.823e-01 | 3.043e-01 | 2.035e-01 |
| 93 | 3.145e-02 | 3.879e-02 | 5.390e-02 | -5.350e-02 | -7.830e-02 | 1.449e-01 |

### S7 Hyperparameters

**Table S8.**
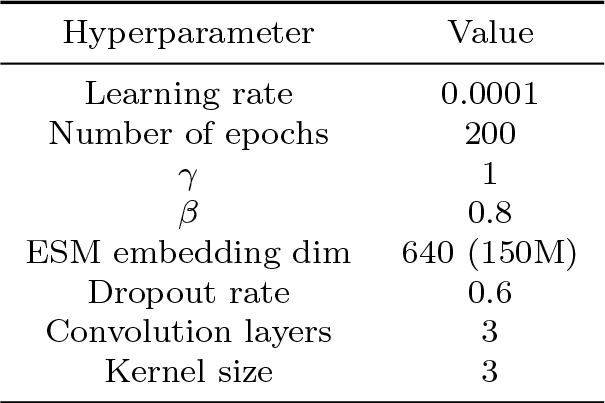
Hyperparameters of the model

| Hyperparameter | Value |
| --- | --- |
| Learning rate | 0.0001 |
| Number of epochs | 200 |
| $\gamma$ | 1 |
| $\beta$ | 0.8 |
| ESM embedding dim | 640 (150M) |
| Dropout rate | 0.6 |
| Convolution layers | 3 |
| Kernel size | 3 |

